# Habitat protection and restoration: win-win opportunities for migratory birds in the Northern Andes

**DOI:** 10.1101/2022.06.01.494397

**Authors:** Ana M. Gonzalez, Nestor Espejo, Dolors Armenteras, Keith A. Hobson, Kevin J. Kardynal, Greg W. Mitchell, Nancy Mahony, Christine A. Bishop, Pablo J. Negret, Scott Wilson

## Abstract

Identifying strategies that offer co-benefits for biodiversity protection, forest restoration and human well-being are important for successful conservation outcomes. In this study, we identified opportunities where forest restoration and rehabilitation programs in Colombia also align with priority areas for the conservation of Neotropical migratory birds. We used citizen science eBird-based abundance estimates to define regions with the highest richness of Neotropical migratory birds of conservation concern at montane elevations in Colombia and aligned these high richness areas with domestic initiatives for forest protection (Forest Areas), restoration (Restoration Areas) and rehabilitation (Rehabilitation Areas). We quantified the location and amounts of these three areas as well as the type of land protection and designation within them, specifically, National Protected Areas, Indigenous Reserves, Afro-descendent territories, and regions affected by poverty and violence that are prioritized for rural development by the Colombian government in Post-conflict Territorially Focused Development Programs (PDET). Almost half of Forest Areas overlapped with PDETs where goals for economic development present a risk of forest loss if not done sustainably. There was a 20% overlap between Forest Areas and Afro-descendant territories and indigenous reserves; most of this overlap was outside of established protected areas thus presenting an opportunity for community forest conservation that benefits migratory birds. We found an alignment of less than 6% between migrant bird focal areas and the priority Restoration and Rehabilitation Planning Areas identified by the Colombian National Restoration Plan indicating less opportunity for these programs to simultaneously benefit Neotropical migrant species. Our approach highlights that timely and efficient conservation of declining migrants depends on identifying the regions and strategies that incorporate local communities as part of the solution to forest loss and degradation in Colombia.

**Highlights:** Colombia covers over half of key wintering areas for migratory birds in South America

Most of the migrants’ overwinter range overlaps with working landscapes

Priority national restoration/rehabilitation areas are ineffective to benefit migrants

Forest conservation needs actions involving vulnerable and minority groups

## Introduction

With multiple competing demands for land use and limited resources for conservation, national institutions mandated to recover declining species are increasingly being tasked to do more with less (Murdoch et al., 2007). Under such circumstances, the success of conservation programs could be enhanced if multiple initiatives are addressed simultaneously, resulting in a win-win approach (López-Cubillos et al., 2022). For example, institutions might differentially focus on either conserving tropical biodiversity or enhancing rural economic development, but both benefit from forest conservation and/or restoration to create a more sustainable landscape for biodiversity and humans alike (Chazdon, 2008). In this study, we show the potential to align conservation efforts to protect declining migratory bird species with forest protection and restoration initiatives in Colombia that aim to improve the welfare and resilience of local communities.

The South American Andes are an important non-breeding (i.e., wintering) area for many Neotropical migratory birds that migrate between their temperate breeding grounds in North America and their wintering grounds in the Neotropics. Among all Neotropical migrant birds, population declines are particularly severe for those species that overwinter in South America where 40% of the total bird abundance has been lost in the last 50 years and 76% of species are in decline (Rosenberg et al., 2019). The northern Andes of Colombia in particular is a region of high conservation importance due to the number of declining Neotropical migrant species occurring there (Wilson et al., 2019) with evidence indicating that habitat loss and degradation in the Andes being the primary cause of these declines (González et al., 2017; Kramer et al., 2018; Wilson et al., 2019, 2018). Recognition of the importance of the Andes to Neotropical migrants has resulted in governmental and non-governmental organizations (NGOs) from Canada and the United States directing resources to conservation efforts in South America, particularly towards those species on federal or regional conservation priority lists where there is a mandate for their recovery (ESA, 1973; SARA, 2002; Wilson et al., 2022).

The Colombian wintering range of most Neotropical migrants of conservation concern falls within mid elevations of mountainous regions between 1000-2300 m asl (Cespedes et al., 2021; Céspedes and Bayly, 2019; Colorado et al., 2012). These regions have had a long and persistent history of high anthropogenic impact with the result that they are now highly modified (Armenteras et al., 2011; Correa Ayram et al., 2020). Across mid elevations along inter-Andean valleys where the core of the human colonization frontier was located, the expansion of crop agriculture and cattle pastures were the major drivers of deforestation (Etter et al., 2008). As a result of these transformations, many montane regions in Colombia are already highly developed, with deforestation hotspots now progressing into more remote and inaccessible regions where remaining native forest is concentrated (Armenteras et al., 2011, 2003; González et al., 2018).

Colombia also went through over 50 years of armed conflict (1964-2016) with the Revolutionary Armed Forces of Colombia (FARC-EP) until a Peace Agreement was signed in 2016 (JEP 2016). The internal conflict further shaped rural landscapes by affecting livelihoods and agricultural production, with small farmers being the most affected (Arias et al., 2014). For instance, between 2000 and 2016 there was some recovery of woody vegetation in the highly deforested 1000-1500 m elevation belt where, driven by forced displacement of the rural population during the armed conflict, the abandonment of pasture and crops (Aide et al., 2019). However, that succession is already being reversed and further forest loss is projected across the country with the implementation of the Colombian post-conflict land reform which incentivizes rural economic development (Negret et al., 2017; Zúñiga-Upegui et al., 2019).

At the same time, concern over forest loss and degradation has led to a growing recognition of the importance of landscape protection and restoration within Colombia (Minambiente, 2015; Ministerio de Agricultura y Desarrollo Rural, 2020), specially for forest dependent birds (Negret et al., 2021). Landscape restoration in particular is emphasized as part of the Ecological Restoration, Rehabilitation and Recuperation of Degraded Areas National Plan (hereafter National Restoration Plan, Vanegas Pinzón et al., 2015). This plan aims to guide and promote integral ecological restoration processes during the next 20 years to recover the structure, composition and function of ecosystems in order to recover biodiversity and ecosystem services in key degraded areas of the country. The plan identified priority regions where the ecosystem can either be i) restored and returned to a natural state when the amount of degradation is low, ii) rehabilitated when returning to the natural state is possible but challenging under moderate levels of degradation, or iii) recuperated to some extent when degradation is severe (Vanegas Pinzón et al., 2015). To examine the potential for dual benefits from this plan to migratory animals and rural human populations, we combined information on the wintering distribution of Neotropical migrant birds with areas targeted by the plan. Importantly, Colombia’s efforts towards increasing protected areas and restoration may present an opportunity to align and coordinate the conservation of Neotropical migrants that are dependent on forested areas and whose recovery will benefit from restoration.

We used data on avian distribution and abundance from the eBird Status and Trends project (Fink et al., 2018) to define priority overwintering regions in the Colombian Andes for six Neotropical migrant species that are the focus of ongoing local and international conservation efforts. We then overlaid focal areas of high richness for these species with the distribution of protected and unprotected forests (hereafter ‘Forest Areas’), and lands prioritized for rehabilitation (‘Rehabilitation Planning Areas’) and restoration (‘Restoration Planning Areas’) in the Colombian National Restoration Plan (Vanegas Pinzón et al., 2015). This was done to delineate areas where forest protection, rehabilitation and restoration has an opportunity to benefit migratory bird conservation. After identifying these Forest, Rehabilitation and Restoration Planning Areas, we quantified the extent of lands stewarded by local communities because we assume this will further influence the likely activities on those lands and the strategies needed to recover migratory birds. To examine stewardship, we identified the overlap between these three areas and 1) Protected Areas; 2) Afro-descendant territories, Indigenous Reserves; 3) Post-conflict Territories, which are regions particularly affected by poverty, violence and inequality prioritized for rural development by the Colombian Government as detailed in Post-conflict Development Programs; and 4) with the jurisdiction of state entities responsible for environmental planning and administrating natural resources at a regional level. Although we focused on Colombia, this approach is also relevant to other countries in northern South America, Central America and the Caribbean developing plans for conservation and restoration of habitat for Neotropical migrants.

## Methods

### Study area

The area of geographic review was across the Andean mountains of Colombia. This region is a global hotspot of threatened and endemic bird species (Orme et al., 2005); and is considered critical for global biodiversity conservation (Myers et al., 2000). The Andes of Colombia are divided into three cordilleras separated by dry and deep basins. The Cauca basin separates the West and Central cordilleras, and the Magdalena basin separates the Central and East cordilleras. Andean forest zoning along the slopes of each cordillera is mainly defined by gradients of temperature and precipitation relative to elevation as follows (Holdridge, 1947): Tropical low-elevation forest (<900–1,000 m), Premontane forest (1000-2000 m asl), Lower Montane forest (2000-3000 m asl), and Montane forest (3000-4000 m asl). Precipitation varies within each elevation belt resulting in further classification of forest into moist forest (1000-2000 mm/year), wet forest (2000-4000 mm/year), and rain forest (>4000 mm/year).

Our study region encompassed the elevation belt between 1000-2300 m asl (hereafter Montane Forest or Montane elevations). We chose this elevation belt for four reasons: 1) Several Neotropical migrants showing population declines depend on forest and agroecosystems located across that range in South America (Céspedes et al., 2021; Céspedes and Bayly, 2019; Colorado et al., 2012; González et al., 2020b; Kramer et al., 2018), 2) the range has a persistent history of loss of natural habitat driven by pastoral, agricultural and urban development that continues today across South America (Armenteras et al., 2011; Tejedor-Garavito et al., 2012), 3) drivers of habitat loss here differ from regions located below 1000 m asl (González et al., 2018, Supplemental Material Table 1), and 4) this range is the geographic focus of ongoing efforts for the conservation of declining Neotropical migrants (Partners In Flight, 2019).

**Table 1.** Land classes used in the analysis. Forest, Restoration and Rehabilitation Planning Areas were defined within Migrant Focal Areas. We assessed the overlap of Protected Areas, Afro-descendant territories and Indigenous Reserves, Post-conflict territories, and Autonomous Regional Corporations (CARs) with Forest, Restoration and Rehabilitation Planning Areas.

| Land classes | Description |
| --- | --- |
| Migrant Focal Areas | Elevation belt from 1000 - 2300 m where 4 or more of the six species are present. |
| Forest Areas | Areas with forest cover, derived from C-LULC. |
| Restoration Planning Areas<br>Rehabilitation Planning Areas | Areas prioritized by the Colombian Restoration Plan. |
| Protected Areas | All Colombian National Protected Areas. |
| Afro-descendant territories and Indigenous Reserves | Lands under a territorial management model. |
| Post-conflict territories (PDETs) | Regions severely affected by poverty, violence and inequality, and prioritized for rural development by the Colombian Government. |
| Autonomous Regional Corporations (CARs) | Entities responsible for environmental planning and administrating natural resources at a regional level. |

### Analysis

Our prioritization targeted six Neotropical migratory species of conservation concern listed on Canada’s Species at Risk Act (SARA 2002) and/or the United States Endangered Species Act (ESA, 1973): Olive-sided Flycatcher (*Contopus cooperi*), Eastern Wood-Pewee (*Contopus virens*), Acadian Flycatcher (*Empidonax virescens*), Golden-winged Warbler (*Vermivora chrysoptera*), Cerulean Warbler (*Setophaga cerulea*), and Canada Warbler (*Cardellina canadensis*). We selected those species because they spend the non-breeding period exclusively in montane areas of Latin America, and had information from eBird on their non-breeding distributions and abundance.

We used a digital elevation model with 900 m resolution to estimate the area in the elevation belt of interest (1000-2300 m asl) in Colombia, Venezuela, Ecuador, and northern Peru and to assess the relative importance of our study area in terms of the total amount of area available as potential habitat for Neotropical migrants.

We used species-specific weekly estimates of relative abundance from the eBird Status and Trends project (Fink et al. 2018) to define the Colombian non-breeding range for the six species, following the approach in (Wilson et al., 2022). The eBird relative abundance estimates are defined as the predicted number of individuals on a one-hour, one-kilometer eBird checklist conducted at the ideal time of day for detection of the species in every week of the year, at a pixel resolution of 2.96 km^2^ (Fink et al., 2018). These relative abundance estimates are generated from an ensemble modeling strategy based on an Adaptive Spatio-Temporal Exploratory Model and include environmental predictors, temporal variation and observer effort to account for detectability (Fink et al., 2020). Thus, for all six species there is a weekly distribution map that includes the estimated relative abundance of the species in each 2.96km^2^ pixel. Using these maps, we focused on the non-breeding season, which was defined from 1 November to 31 March and, then we estimated the average relative abundance per pixel across all weeks for each species. Migratory species often have low abundance at range edges and we only selected pixels that represented a cumulative 95% of total abundance of each species to focus on their core nonbreeding range. The rasters for the 95% of total abundance for each species were then converted to a presence-absence raster where any pixel that contributed to the 95% abundance was assigned a 1 and all other pixels were assigned a 0. These presence-absence rasters for the six species were then stacked using package Raster (Hijmans, 2019) in R version 4.0.3 (R Core Team, 2020) and the stacked estimates for each pixel were used to estimate species richness per pixel (i.e. the number of focal species present in each pixel). From this derived map of species richness, we then selected areas with 4 or more species present to focus our analysis on regions that provide benefit for multiple migratory species. As a final step, we restricted this derived species richness layer to the 1000 to 2300 m asl range within Colombia and defined this as our Migrant Focal Area. We acknowledge that a richness-based distribution map will not allow us to incorporate differences in abundance into the consideration of priority areas for migratory birds. However, the six species differ in abundance across the wintering grounds and we chose a method that treated all six species equally rather than favoring those that are more abundant.

We used the 2018 land use and land cover maps (C-LULC) applying the Corine and Land Cover methodology adapted for Colombia (IDEAM, 2018) to derive Forested Areas within Migrant Focal Areas and to assess other land uses (Supplemental Material Table 2). Forest Areas were considered as land with a minimum tree canopy cover of 30%, minimum canopy height of 5 m, and a minimum area of 1 ha (IDEAM, 2018). Within the Migrant Focal Areas we also identified regions that overlapped with areas prioritized for rehabilitation and restoration in the National Restoration Plan (Vanegas Pinzón et al., 2015); these overlapping areas represented our Restoration and Rehabilitation Planning Areas. Forested, restoration and rehabilitation areas were transformed to a raster format of 100 m resolution/pixel.

After identifying our Forest, Restoration and Rehabilitation Planning Areas, we assessed their protected status and stewardship by examining the area covered by 1) National Protected Areas (RUNAP 2021), 2) unprotected areas, 3) Indigenous Reserves and Afro-descendent territories (IGAC, 2021), 4) regions prioritized in Post-conflict Territorially Focused Development Programs (PDET hereafter post-conflict territories, IGAC 2021), and 5) Autonomous Regional Corporations (CARs), which are the state entities responsible for decentralized environmental governance by implementing and enforcing law and policies developed by the Ministry of the Environment (Table 1). As such, the 33 CARs across the country are responsible for regional environmental planning and administrating natural resources within their jurisdictions. The total area and percentage of overlap of the Migrant Focal Areas with the land classes mentioned above was calculated using ArcMap 10.5 (ESRI, 2019).

National Protected Areas were classified according to their management objectives as follows (IUCN, 2008, Supplemental Material Table 3): (Ib) Wilderness Areas which are largely unmodified and their primary objective is to preserve their natural condition; (II) National Parks which are large natural areas set apart to protect their natural biodiversity, ecological structure and promote education and recreation; (V) Protected Landscapes which are areas that maintain their integrity by a balanced interaction between people and nature through traditional management practices; and (VI) Protected Areas with Sustainable Use of Natural Resources where most of the area is in a natural condition, but some areas are under sustainable natural resource management. All protected areas form the National Protected Areas System (SINAP).

## Results

The elevation belt between 1000 and 2300 m across Colombia, Venezuela, Ecuador and northern Peru covers 22,575,202 ha. Most of this elevation range falls in Colombia (57%) followed by Ecuador (18%), Peru (13%), and Venezuela (11%). Migrant Focal Areas within that elevation belt covered 10,831,472 ha across the three Andean chains in Colombia. Forest Areas covered 4,115,368 ha corresponding to 38% of Migrant Focal Areas. Most of the non-forest areas were covered by crop (30%), followed by pasture (17%), early successional habitats (12%), and agroforestry systems (3%, Supplemental Material Table 2).

### Forest Areas

Forest Areas were concentrated in the western slope of the West Andes and the eastern slope of the East Andes (Figure 1). Post-conflict territories had the largest amount of land overlapping with Forest Areas (47%), followed by National Protected Areas (30%), and Afro-descendant territories and Indigenous Reserves (19%) (Figure 2a, 2b). Only 2% of Forest Areas in Afro-descendant territories and indigenous reserves overlapped with protected areas. Protected areas within our Forest Area category were primarily represented by National Parks (IUCN category II, 79% coverage) followed by Protected Areas with Sustainable Use of Natural Resources (IUCN category VI, 20% coverage).

**Figure 1.**
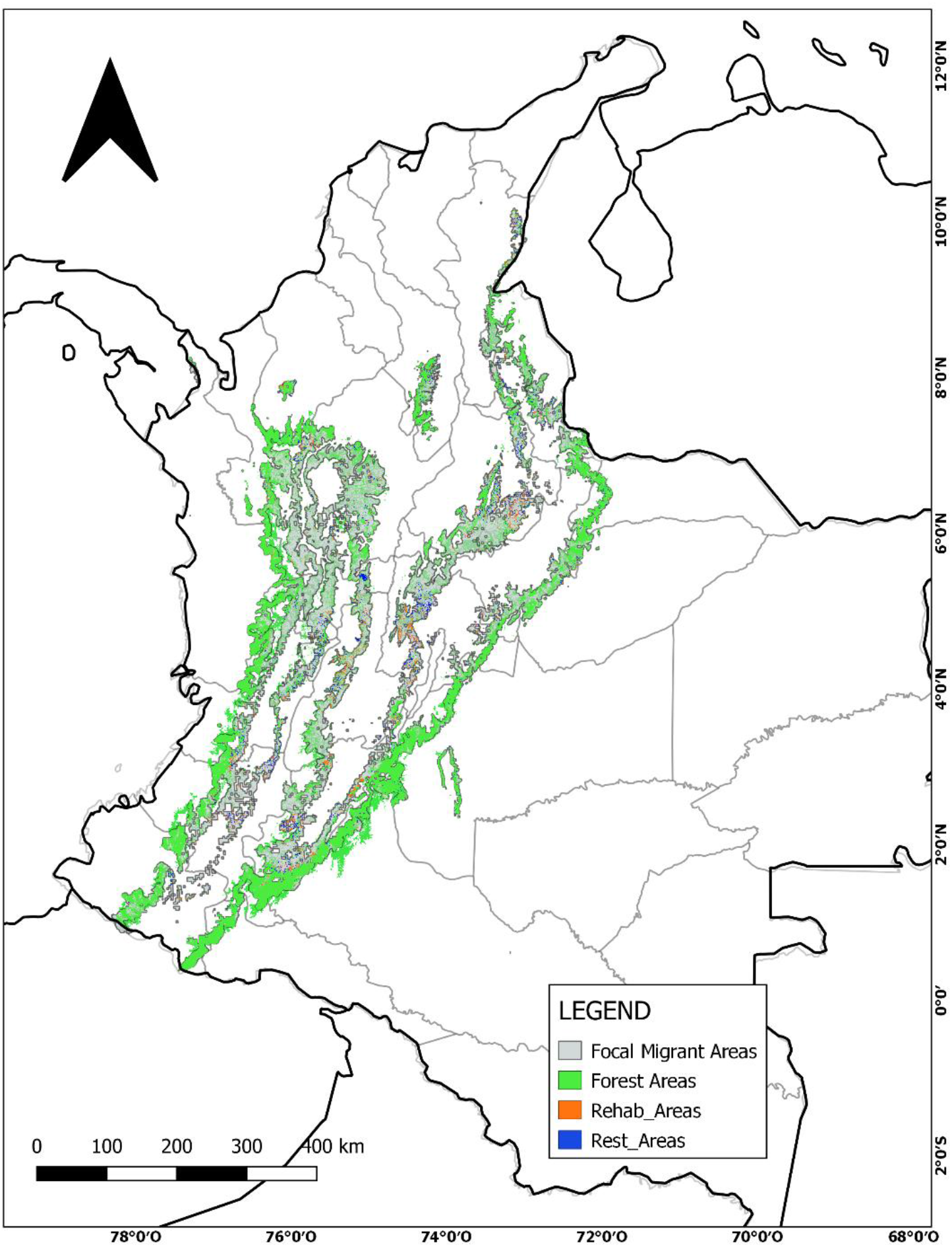
Migrant Focal Areas: Elevation belt from 1000 - 2300 m asl in Colombia where four or more of the six focal declining species are present. We identified and defined Forest, Rehabilitation and Restoration Planning Areas within Migrant Focal Areas. Gray regions indicate degraded areas within Migrant Focal Areas that are not prioritized by the Colombian National Restoration Plan.

**Figure 2.**
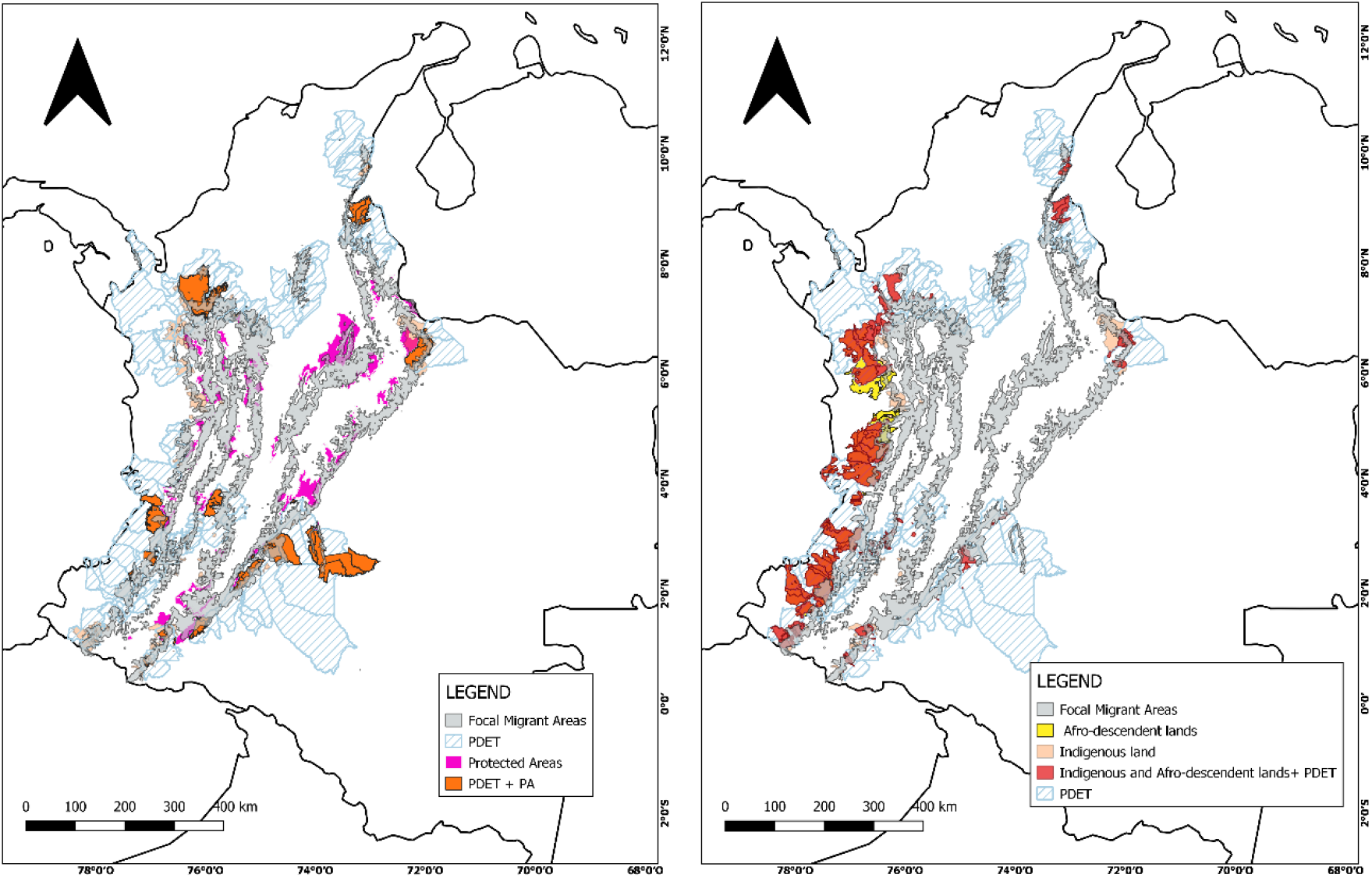
Migrant Focal Areas: Elevation belt from 1000 - 2300 m asl where four or more of the six focal declining species are present. A. Overlap of Migrant Focal Areas with Post-conflict territories (PDET), Protected Areas (PA), and overlap between PDET and PA. B. Overlap of Migrant Focal Areas with Afro-descendent lands, indigenous lands, and overlap between Indigenous and Afro-descendent lands and PDET.

Forest Areas overlapped with National Parks primarily in the eastern slope of the East Andes including Cordillera de Los Picachos, Cocuy, Serrania de Los Chrumbelos and Serrania de los Yariguies; and the western slope of the West Andes including Paramillo, Sierra de la Macarena and Farallones de Cali National Parks (Figure 2a). Over 50% of Forest Areas fell within the jurisdiction of five CARs localized in the east and southwest of the country: Corpoamazonia, Corponariño, Corporación Autónoma Regional del Cauca, Codechoco, Corporinoquia, and Cormacarena (Supplemental Material Table 4, and Figure 2).

### Restoration and Rehabilitation Planning Areas

The overlap between Migrant Focal Areas and areas prioritized by the Colombian National Restoration Plan was low. Specifically, Restoration and Rehabilitation Planning Areas covered only 2.5% and 2.9% of Migrant Focal Areas respectively (Figure 1). Most Restoration Planning Areas were located in the Magdalena valley while Rehabilitation Planning Areas were located in the Cauca Valley, along the western slope of the Central Andes (Figure 1). Within the Restoration and Rehabilitation Planning Areas,32% and 19% of the area was protected, respectively. However, unlike Forest Areas, where protected areas were primarily represented by National Parks (IUCN class II), the majority of protected area falling within Restoration and Rehabilitation Planning Areas was designated as Sustainable Use of Natural Resources (IUCN class VI). Specifically, 55% and 67% of the protected area within Restoration and Rehabilitation Planning Areas, respectively, fell within this class.

Afro-descendant territories and Indigenous Reserves overlapped with less than 3% of both Restoration and Rehabilitation Planning Areas. There was a greater representation of post-conflict territories, which represented 19% and 16% of Restoration and Rehabilitation Planning Areas, respectively. The CARs of Cauca, Tolima, Santander, Cundinamarca and Huila located in the Central and East Andes had over 50% of Restoration and Rehabilitation Planning Areas within their jurisdictions (Supplemental Material Table 4 and Figure 2).

## Discussion

The scale and complexity of conservation issues faced by Neotropical migrants during the wintering period requires the identification and targeting of regions across remaining forested habitats and disturbed landscapes used by migrants as well as the collaboration of individuals across administrative boundaries, land ownership and political jurisdictions (Scarlett and McKinney, 2016; Schuster et al., 2019). Colombia is a critically important wintering area for numerous Neotropical migrant birds in decline (Cespedes et al., 2021; González et al., 2017; Wilson et al., 2018) and is also a global biodiversity hotspot (Myers et al., 2000). By focusing this study on Colombia, we aimed to provide information on the extent to which goals for migratory bird conservation and sustainable development overlap within the country to aid future planning but also to provide a framework by which a similar process could be applied to other Latin American countries that are of high importance for migratory species. Our study highlighted four key results regarding the degree of alignment between the distributions of Neotropical migrants of conservation concern and the ongoing programs within Colombia to protect or recover forests. First, most of the Forest Areas fall along the west slope of the western Andes and the east slope of the Eastern Andes with very little representation in central Andean areas. Second, almost half of Forested Areas overlapped with post-conflict territories where goals for economic development present a risk of forest loss if not done sustainably. Third, there was a large overlap between Forested Areas and Afro-descendant territories and Indigenous Reserves; most of this overlap was outside of established protected areas thus presenting an opportunity for community forest conservation that potentially benefits migratory birds. Fourth, we found very limited alignment between migrant focal areas and the priority Restoration and Rehabilitation Planning Areas identified by the Colombian National Restoration Plan; this lack of alignment suggests that alternative conservation strategies will be needed for migratory birds in these areas.

Collaborative efforts between governments, NGOs and academia in North and Latin America have allowed the development of conservation plans to address threats for species such as Canada Warbler (ECCC and BirdLife International, 2021), Golden-winged Warbler (Bennett et al., 2018), and Cerulean Warbler (Fundación ProAves et al., 2010) during the overwintering period. Ongoing efforts such as the development of the Central and South America mid-elevation forest Conservation Investment Strategy (Partners In Flight, 2019) will integrate the strategies and actions of those three plans, and will benefit other Neotropical migrants and resident species across montane elevations in Central and South America. Despite those planning efforts to conserve migratory birds at montane elevations and the recognition of the importance of full-annual cycle conservation approaches to reverse declines, efforts directed from North America are still limited by current legislation. For instance, Canadian legislation such as the Migratory Bird Conservation Act focuses protection only on the breeding grounds and the Species at Risk Act only identifies critical habitat within Canada. Efforts within Canada alone are not enough to conserve Neotropical migrants as evidenced by the ongoing decline of 80% of the species breeding in the boreal or hardwood forest in Canada and overwintering in the Colombian Andes (Hobson and Wilson, 2020). Additional programs and agreements that allow for conservation investments outside North America such as those already undertaken in the United States to conserve birds in the Caribbean and Latin America through the Neotropical Migratory Bird Conservation Act (NMBCA; Public Law 106-247) are urgently needed to implement conservation strategies in critical areas such as the Andes of Northern South America.

Forest cover along the West and East Andes presents opportunities for conservation efforts including the declaration of new protected areas but clearly necessitates additional strategies across private lands and in more central regions where little forest cover remains. For instance, Forest Areas, including Afro-descendant territories and Indigenous lands, have high potential for the implementation of community-based conservation strategies such as ecotourism. Indeed 20% of Forest Areas lay within the departments of Cauca and Nariño in the West Andes which have the highest diversity of birds in the country (Vélez et al., 2021). The potential economic and conservation benefits of bird-watching have also been highlighted in regions prioritized for economic development such as post-conflict territories (Ocampo-Peñuela and Winton, 2017) which covered 50% of Forest Areas. Support for bird-watching tourism including infrastructure investment, training of local guides, and promoting those regions as world-class birding destinations is needed to meet goals for economic growth while protecting forest (Ocampo-Peñuela and Winton, 2017, Echeverry et al. 2022).

The large overlap between Forest Areas and post-conflict territories and the emphasis on economic development in those regions presents a major challenge to the conservation of migratory species dependent on forest but also presents an important opportunity to incorporate needs of migrant birds into planning efforts. Such efforts have the potential to jointly benefit wildlife conservation and sustainable human livelihoods. A key aspect of the Colombian Peace Agreement is the need for an integral rural reform in order to address the historical drivers of persistent violence and armed conflict such as the concentration of land ownership and income, and farmer displacement (JEP, 2016). Post-conflict Development Programs are the planning and management instruments to implement the components of the rural reform and are the product of a consensus between local authorities and civil population with the aim of identifying regional priority needs and propose remediation projects (De la Rosa and Contreras, 2018).

Community participation, the revitalization of community-based organizational processes in the territory, and the alignment of interinstitutional efforts have been key for the implementation of Post-conflict Development Programs across several regions in Colombia (Barbosa et al., 2021). Those tools have allowed farmers and Indigenous and Afro-descendant communities to take advantage of the planning spaces of the Post-conflict Development Programs to strengthen their organizational and productive processes. One of the rural reform critical points was the Environmental Zoning Plan across post-conflict territories which defines management and conservation of areas of special environmental interest and facilitates the allocation of governmental support for community-based conservation programs including payment for environmental services and support to sustainable food production systems. For instance in October 2021, the Colombian government approved $22 million USD for environmental projects across post-conflict territories (Presidencia de la Republica de Colombia, 2021). We urge national and international organizations to approach local organizations across post-conflict territories and align management actions for the conservation of migratory birds and other threatened species (e.g., Wilson et al. 2022) with regional planning and ongoing conservation strategies. Aligning conservation efforts for Neotropical migrants with post-conflict territories would affect half of the area of Forested Areas and would support sustainable social and environmental rural development. The need to align conservation efforts also applies to other regions across Migrant Focal Areas where engaging with regional or local institutions such as CARs is needed to maximize financial and logistic resources and to design and implement projects according to regional and local planning needs.

Despite positive outcomes in some post-conflict territories, rural communities still face several challenges for environmental peacebuilding. Poor government commitment to implement the Peace Agreement and weak state presence and enforcement of environmental polices have resulted in increased deforestation rates, homicides, threats against social-environmental leaders, and pose unique socio-politic challenges for biodiversity conservation and management across many regions (Clerici et al., 2020; Graser et al., 2020; Negret et al., 2019; Prem et al., 2020). At a regional level, integrated management of natural resources between the civil society, international corporations, and national environmental agencies is urgently needed to implement initiatives that contribute to sustainable peace including the adoption of sustainable rural development and the empowerment of local communities (Graser et al., 2020; Torres Rodríguez et al., 2020).

Almost 20% of remaining Forest Areas were located within Afro-descendant territories and Indigenous Reserves mainly in the West Andes however only 3% of the area is protected. Implementing conservation, or restoration, would be impossible across several regions without the leadership of Indigenous and other local communities and without recognizing their land rights. Engaging effectively with those communities to define conservation objectives in the context of economic development opportunities is needed to mitigate deforestation in these remote regions prone to deforestation (Negret et al., 2017). For example, this could be done through programs such as the conservation business strategies (Partners In Flight, 2019), where North American organizations involved in migratory bird conservation have an opportunity to align with local conservation groups that work with local communities to reduce deforestation and engage in restoration.

Established Protected areas covered about 30% of Forest Areas indicating that most remaining forested areas important for migratory birds lack any formal protection. This pattern is consistent with the poor protection of migratory birds in the global protected areas system (Runge et al., 2015). Our results show where the differing levels of protection afforded the Forest Areas might benefit from effective management and additional protection. Lack of effective management across protected areas represents a threat for declining Neotropical migrants and biodiversity (Runge et al., 2015). For instance, in Colombia, many protected areas had been found ineffective preventing forest loss (Negret et al., 2020). Moreover deforestation in protected areas and surrounding buffer areas increased after the Peace Agreement due, in part, to historical financial and operational weakness of the national government to enforce effective protection of public conservation areas (Clerici et al., 2020). The diversity of management strategies within the Colombian National Protected Areas System offers an opportunity for conservation planning across diverse landscapes. For instance, Protected Areas with Sustainable Use of Natural Resources such as Civil Society Natural Reserves would benefit declining Neotropical migrants through the conservation or restoration of forest in private lands. This approach can be implemented in highly deforested regions such as the Cauca and Magdalena Valley or in remote regions with extensive remaining forest where economic development is expected, such as the West Andes.

The poor alignment between Migrant Focal Areas and Restoration and Rehabilitation Planning Areas is largely explained by the methodology used in the National Restoration Plan to identify key restoration regions. Areas susceptible to restoration and rehabilitation were identified, in part, by assessing the type of land-use change between the periods 2000-2002 and 2005-2009, and by identifying the regions affected by deforestation during four periods between 1990-2012 (Vanegas Pinzón et al., 2015). However, the Colombian Andes have a persistent history of high human impact since pre-Columbian times and have experienced continuous population growth and economic development since the 1970’s (Correa Ayram et al., 2020; Etter et al., 2008; Rodríguez Eraso et al., 2016). The vast majority of Migrant Focal Areas experienced forest loss prior to the 1990-2012 period used in the National Plan and therefore were not selected as areas for restoration and rehabilitation; thus, new approaches to target key restoration areas to benefit Neotropical migrants are needed. Indeed, over 60% of Migrant Focal Areas are covered by non-forested habitats and these are primarily productive lands for crops, pastures, and agroforestry systems. Increasing habitat availability and suitability for Neotropical migrants across those regions thus may largely depend on implementing conservation approaches within working lands that simultaneously support productive landscapes and human well-being while maintaining biodiversity and ecosystem services (Kremen and Merenlender, 2018). For instance, the implementation of biodiversity-based management techniques such as agroforestry and silvopastural systems would increase the resilience of crop production to climate change (Vaast et al., 2016), and enhance the livelihood and food security of farmers (Hernandez-Aguilera et al., 2019; Waldron et al., 2017) while providing suitable habitat for Neotropical migrants (Colorado et al., 2018; González et al., 2020a, 2020b; McDermott et al., 2015).

### Conclusions

We provide Colombian and international conservation agencies with information needed to plan and implement avian conservation initiatives that can overlap with the socio-political, cultural and ethnical local context within Colombia. We also recognize that attempts to address conservation without the direct involvement and leadership of minority rural communities will not only continue to be unethical, but will also likely result in unsuccessful conservation outcomes (Artelle et al., 2019). Extreme poverty in rural areas of Colombia is over three times as high as in urban areas (DANE, 2017), and higher levels of poverty among peasants, Indigenous and Afro-Colombian people are largely related to inequity in the distribution of land tenure which in turns increases deforestation pressure (Armenteras et al., 2019). The level of land tenure security can hinder the capacity of conservation organizations to influence land management decisions (Robinson et al., 2018). Promoting the legal recognition and protection of land and territorial rights of indigenous, Afro-Colombians, and rural communities, including their rights to self-governance, is key for the enrollment of those communities in sustainable conservation programs that require tenure security such as Payment for Ecosystem Services (PES) and to achieve effective conservation (Robinson et al., 2018; Worsdell et al., 2020).

Local communities are an integral part of montane ecosystems used by declining Neotropical migrants, and timely and efficient conservation depends on identifying the regions and strategies that incorporate people as part of the solution to habitat loss and degradation (Armsworth et al., 2007; Dayer et al., 2020). Indeed, conservation approaches that support economical development and human wellbeing such as PES and integrated landscape management are often prioritized by conservation agencies in Latin America (Doak et al., 2014). Although these approaches have resulted in successful institutional planning and coordination, on-the-ground tangible outcomes in agriculture, livelihoods, and conservation domains are scarce. Some of the challenges that we need to address include unsupportive policies, lack of engagement of key stakeholders such as government and private sector and poor continuous financial and technical support to allow for adaptation to new frameworks (Estrada-Carmona et al., 2014). Moving beyond conservation planning and building strong partnerships to implementing on-the-ground strategies is a priority to produce tangible out comes that benefit humans and migratory and resident species together in the Neotropics.

## Acknowledgments

Funding for this study was provided by Environment and Climate Change Canada – Science and Technology Branch. We are grateful to Nicholas J. Bayly for his feedback on the final draft of the manuscript.

## References

[ESA] Endangered Species Act, 1973. Endangered Species Act of 1973 (16 U.S.C. 1531–1544, 87 Stat. 884).

Aide, T.M., Grau, H.R., Graesser, J., Andrade-Nuñez, M.J., Aráoz, E., Barros, A.P., Campos-Cerqueira, M., Chacon-Moreno, E., Cuesta, F., Espinoza, R., Peralvo, M., Polk, M.H., Rueda, X., Sanchez, A., Young, K.R., Zarbá, L., Zimmerer, K.S., 2019. Woody vegetation dynamics in the tropical and subtropical Andes from 2001 to 2014: Satellite image interpretation and expert validation. Glob. Chang. Biol. 25, 2112–2126. https://doi.org/10.1111/gcb.14618

Arias, M.A., Ibañez, A.M., Zambrano, A., 2014. Agricultural Production Amid Conflict: The Effects of Shocks, Uncertainty, and Governance of Non-State Armed Actors Author & abstract. Bogotá D.C., Colombia.

Armenteras, D., Gast, F., Villareal, H., 2003. Andean forest fragmentation and the representativeness of protected natural areas in the eastern Andes, Colombia. Biol. Conserv. 113, 245–256. https://doi.org/10.1016/S0006-3207(02)00359-2

Armenteras, D., Negret, P., Melgarejo, L.F., Lakes, T.M., Londoño, M.C., García, J., Krueger, T., Baumann, M., Davalos, L.M., 2019. Curb land grabbing to save the Amazon. Nat. Ecol. Evol. https://doi.org/10.1038/s41559-019-1020-1

Armenteras, D., Rodríguez, N., Retana, J., Morales, M., 2011. Understanding deforestation in montane and lowland forests of the Colombian Andes. Reg. Environ. Chang. 11, 693–705. https://doi.org/10.1007/s10113-010-0200-y

Armsworth, P.R., Chan, K.M.A., Daily, G.C., Ehrlich, P.R., Kremen, C., Ricketts, T.H., Sanjayan, M.A., 2007. Ecosystem-service science and the way forward for conservation. Conserv. Biol. https://doi.org/10.1111/j.1523-1739.2007.00821.x

Artelle, K.A., Zurba, M., Bhattacharrya, J., Chan, D.E., Brown, K., Housty, J., Moola, F., 2019. Supporting resurgent Indigenous-led governance: A nascent mechanism for just and effective conservation. Biol. Conserv. 240. https://doi.org/10.1016/j.biocon.2019.108284

Barbosa, M. del P., Garcia, A., Maya, M., Sanabria, O., Valderrama, M., 2021. Logros y aciertos en la implementación de los Programas de Desarrollo con Enfoque Territorial – PDET.

Bennett, R.E., Rothman, A., Rosenberg, K. V, Rodríguez, F., 2018. Plan de conservación de la Reinita Alidorada (Vermivora chrysoptera) para la temporada no reproductiva. Golden-winged Warbler working group.

Cespedes, L., Wilson, S., Bayly, N.J., 2021. Community modeling reveals the importance of elevation and land cover in shaping migratory bird abundance in the Andes. Ecol. Appl. e02481. https://doi.org/doi.org/10.1002/eap.2481

Céspedes, L.N., Bayly, N.J., 2019. Over-winter ecology and relative density of Canada Warbler Cardellina canadensis in Colombia: The basis for defining conservation priorities for a sharply declining long-distance migrant. Bird Conserv. Int. 29, 232–248. https://doi.org/10.1017/S0959270918000229

Céspedes, L.N., Wilson, S., Bayly, N.J., 2021. Community modeling reveals the importance of elevation and land cover in shaping migratory bird abundance in the Andes. Ecol. Appl. 31, e02481.

Chazdon, R.L., 2008. Beyond deforestation: Restoring forests and ecosystem services on degraded lands. Science (80-.). 320, 1458–1460. https://doi.org/10.1126/science.1155365

Clerici, N., Armenteras, D., Kareiva, P., Botero, R., Ramírez-Delgado, J.P., Forero-Medina, G., Ochoa, J., Pedraza, C., Schneider, L., Lora, C., Gómez, C., Linares, M., Hirashiki, C., Biggs, D., 2020. Deforestation in Colombian protected areas increased during post-conflict periods. Sci. Rep. 10. https://doi.org/10.1038/s41598-020-61861-y

Colorado, G.J., Hamel, P.B., Rodewald, A.D., Mehlman, D., 2012. Advancing our understanding of the non-breeding distribution of Cerulean Warbler (Setophaga cerulea) in the Andes. Ornitol. Neotrop. 23, 307–315.

Colorado Z, G.J., Mehlman, D., Valencia-C, G., 2018. Effects of floristic and structural features of shade agroforestry plantations on the migratory bird community in Colombia. Agrofor. Syst. 92. https://doi.org/10.1007/s10457-016-0034-9

Correa Ayram, C.A., Etter, A., Díaz-Timoté, J., Rodríguez Buriticá, S., Ramírez, W., Corzo, G., 2020. Spatiotemporal evaluation of the human footprint in Colombia: Four decades of anthropic impact in highly biodiverse ecosystems. Ecol. Indic. 117, 106630. https://doi.org/10.1016/j.ecolind.2020.106630

DANE, 2017. Pobreza Monetaria y Multidimensional en Colombia 2017 [WWW Document]. URL https://www.dane.gov.co/index.php/estadisticas-por-tema/pobreza-y-condiciones-de-vida/pobreza-y-desigualdad/pobreza-monetaria-y-multidimensional-en-colombia-2017#pobreza-monetaria-y-multidimensional-en-colombia-2017 (accessed 4.8.21).

Dayer, A.A., Silva-Rodríguez, E.A., Albert, S., Chapman, M., Zukowski, B., Ibarra, J.T., Gifford, G., Echeverri, A., Martínez-Salinas, A., Sepúlveda-Luque, C., 2020. Applying conservation social science to study the human dimensions of Neotropical bird conservation. Condor 122. https://doi.org/10.1093/condor/duaa021

De la Rosa, M.D., Contreras, D., 2018. Instrumentos administrativos para la paz: Programas de Desarrollo con Enfoque Territorial (pdet), in: Lecturas Sobre Derecho de Tierras. Tomo II. Universidad Externado de Colombia, Bogotá, D.C., pp. 273–310.

Doak, D.F., Bakker, V.J., Goldstein, B.E., Hale, B., 2014. What is the future of conservation? Trends Ecol. Evol. https://doi.org/10.1016/j.tree.2013.10.013

ECCC and BirdLife International, 2021. Canada Warbler full-life-cycle Conservation Action Plan.. Gatineau, Québec, Canada and Quito, Ecuador.

Echeverry, A., Smith, J.R., MacArthur-Waltz, D., Lauck, K.S., Anderson, C.B., Vargas, R.M., Quesada, I.A., Wood, S.A., Chaplin-Kramer, R., and Daily, GC. 2022. Biodiversity and infrastructure interact to drive tourism to and within Costa Rica. Proceedings of the National Academy of Sciences. In press.

ESRI, 2019. ArcGIS Desktop: Release 10.

Estrada-Carmona, N., Hart, A.K., DeClerck, F.A.J., Harvey, C.A., Milder, J.C., 2014. Integrated landscape management for agriculture, rural livelihoods, and ecosystem conservation: An assessment of experience from Latin America and the Caribbean. Landsc. Urban Plan. 129, 1–11. https://doi.org/10.1016/j.landurbplan.2014.05.001

Etter, A., McAlpine, C., Possingham, H., 2008. Historical patterns and drivers of landscape change in Colombia since 1500: a regionalized spatial approach. Ann. Assoc. Am. Geogr. 98, 2–23. https://doi.org/10.1080/00045600701733911

Fink, D., Auer, T., Johnston, A., Ruiz-Gutierrez, V., Hochachka, W.M., Kelling, S., 2020. Modeling avian full annual cycle distribution and population trends with citizen science data. Ecol. Appl. 30. https://doi.org/10.1002/eap.2056

Fink, D., Auer, T., Johnston, A., Strimas-Mackey, M., Illif, M., Kelling, S., 2018. eBird Status and Trends, Version November 2018, eBird Status and Trends, Version November 2018. Ithaca, NY, USA.

Fundación ProAves, America Bird Concervancy, El Grupo Cerúleo, 2010. Conservation Plan for the Cerulean Warbler on its nonbreeding range—Plan de conservación para la Reinita Cerúlea sobre su rango no reproductivo. Conserv. Colomb. 12, 1–62.

González, A.M., Bayly, N.J., Colorado, G.J., Hobson, K.A., 2017. Topography of the Andes mountains shapes the wintering distribution of a migratory bird. Divers. Distrib. 23, 118–129. https://doi.org/10.1111/ddi.12515

González, A.M., Bayly, N.J., Hobson, K.A., 2020a. Earlier and slower or later and faster: Spring migration pace linked to departure time in a Neotropical migrant songbird. J. Anim. Ecol. 89, 2840–2851. https://doi.org/10.1111/1365-2656.13359

González, A.M., Wilson, S., Bayly, N.J., Hobson, K.A., 2020b. Contrasting the suitability of shade coffee agriculture and native forest as overwinter habitat for Canada Warbler (Cardellina canadensis) in the Colombian Andes. Condor 122, 1–12. https://doi.org/10.1093/condor/duaa011

González, J., Cubillos, Á., Chadid, M., Cubillos, A., Arias, M., Zúñiga, E., Joubert, F., Pérez, I., Berrío, V., 2018. Caracterización de las principales causas y agentes de la deforestación a nivel nacional período 2005-2015. Bogotá, Colombia.

Graser, M., Bonatti, M., Eufemia, L., Morales, H., Lana, M., Löhr, K., Sieber, S., 2020. Peacebuilding in rural Colombia-a collective perception of the Integrated Rural Reform (IRR) in the department of Caqueta (Amazon). Land 9. https://doi.org/10.3390/land9020036

Hernandez-Aguilera, J.N., Conrad, J.M., Gómez, M.I., Rodewald, A.D., 2019. The Economics and Ecology of Shade-grown Coffee: A Model to Incentivize Shade and Bird Conservation. Ecol. Econ. 159, 110–121. https://doi.org/10.1016/j.ecolecon.2019.01.015

Hijmans, R.J., 2019. raster: Geographic data analysis and modeling.

Hobson, K.A., Wilson, S., 2020. The avian conservation crisis, canada’s international record, and the need for a new path forward. Avian Conserv. Ecol. 15. https://doi.org/10.5751/ACE-01756-150222

Holdridge, L.R., 1947. Determination of world plant formations from simple climatic data. Science (80-.). 105, 367–368. https://doi.org/10.1126/science.105.2727.367

IDEAM - Instituto de Hidrología, M. y E.A., 2018. Cobertura de la Tierra Metodología CORINE Land Cover Adaptada para Colombia Periodo 2018. República de Colombia. Escala 1:100.000. Año 2021 [WWW Document]. URL http://geoservicios.ideam.gov.co/geonetwork/srv/eng/catalog.search#/metadata/285c4d0a-6924-42c6-b4d4-6aef2c1aceb5

IGAC. Instituto geográfico Agustín Codazzi, 2021. Sistema de información geográfica para la planeación y el ordenamiento territorial.

IUCN. International Union for Conservation of Nature, 2008. Guidelines for applying protected area managment categories. Gland, Switzerland.

JEP. Jurisdicción Especial para la paz, 2016. Acuerdo final para la terminación del conflicto y la construcción de la paz estable y duradera [WWW Document]. URL https://www.jep.gov.co/Normativa/Paginas/Acuerdo-Final.aspx (accessed 3.29.21).

Kramer, G.R., Andersen, D.E., Buehler, D.A., Wood, P.B., Peterson, S.M., Lehman, J.A., Aldinger, K.R., Bulluck, L.P., Harding, S., Jones, J.A., Loegering, J.P., Smalling, C., Vallender, R., Streby, H.M., 2018. Population trends in Vermivora warblers are linked to strong migratory connectivity. Proc. Natl. Acad. Sci. U. S. A. 115, E3192–E3200. https://doi.org/10.1073/pnas.1718985115

Kremen, C., Merenlender, A.M., 2018. Landscapes that work for biodiversity and people. Science (80-.). https://doi.org/10.1126/science.aau6020

López-Cubillos, S., Runting, R.K., Suárez-Castro, A.F., Williams, B.A., Armenteras, D., Manuel Ochoa-Quintero, J., McDonald-Madden, E., 2022. Spatial prioritization to achieve the triple bottom line in Payment for ecosystem services design. Ecosyst. Serv. 55. https://doi.org/10.1016/j.ecoser.2022.101424

McDermott, M.E., Rodewald, A.D., Matthews, S.N., 2015. Managing tropical agroforestry for conservation of flocking migratory birds. Agrofor. Syst. 89. https://doi.org/10.1007/s10457-014-9777-3

Minambiente, 2015. Plan nacional de restauración. Restauración ecológica, rehabilitación y recuperación de áreas disturbadas. Bogotá D.C., Colombia.

Ministerio de Agricultura y Desarrollo Rural, 2020. Decreto 130 de 2020. “Por el cual se sustituye el Título 1 de la Parte 3 del Libro 2 del Decreto 1071 de 2015, Decreto Único Reglamentario del Sector Agropecuario, Pesquero y de Desarrollo Rural, relacionado con el Certificado de Incentivo Forestal-CIF.” Bogotá D.C., Colombia.

Murdoch, W., Polasky, S., Wilson, K.A., Possingham, H.P., Kareiva, P., Shaw, R., 2007. Maximizing return on investment in conservation. Biol. Conserv. 139, 375–388. https://doi.org/10.1016/j.biocon.2007.07.011

Myers, N., Mittermeler, R.A., Mittermeler, C.G., Da Fonseca, G.A.B., Kent, J., 2000. Biodiversity hotspots for conservation priorities. Nature. https://doi.org/10.1038/35002501

Negret, P.J., Allan, J., Braczkowski, A., Maron, M., Watson, J.E.M., 2017. Need for conservation planning in postconflict Colombia. Conserv. Biol. 31, 499–500. https://doi.org/10.1111/cobi.12935

Negret, P.J., Marco, M. Di, Sonter, L.J., Rhodes, J., Possingham, H.P., Maron, M., 2020. Effects of spatial autocorrelation and sampling design on estimates of protected area effectiveness. Conserv. Biol. 34. https://doi.org/10.1111/cobi.13522

Negret, P.J., Maron, M., Fuller, R.A., Possingham, H.P., Watson, J.E.M., Simmonds, J.S., 2021. Deforestation and bird habitat loss in Colombia. Biol. Conserv. 257. https://doi.org/10.1016/j.biocon.2021.109044

Negret, P.J., Sonter, L., Watson, J.E.M., Possingham, H.P., Jones, K.R., Suarez, C., Ochoa-Quintero, J.M., Maron, M., 2019. Emerging evidence that armed conflict and coca cultivation influence deforestation patterns. Biol. Conserv. 239, 108176. https://doi.org/10.1016/j.biocon.2019.07.021

Ocampo-Peñuela, N., Winton, R.S., 2017. Economic and Conservation Potential of Bird-Watching Tourism in Postconflict Colombia. Trop. Conserv. Sci. 10. https://doi.org/10.1177/1940082917733862

Orme, C.D.L., Davies, R.G., Burgess, M., Eigenbrod, F., Pickup, N., Olson, V.A., Webster, A.J., Ding, T.S., Rasmussen, P.C., Ridgely, R.S., Stattersfield, A.J., Bennett, P.M., Blackburn, T.M., Gaston, K.J., Owens, I.P.F., 2005. Global hotspots of species richness are not congruent with endemism or threat. Nature 436, 1016–1019. https://doi.org/10.1038/nature03850

Partners In Flight, 2019. Central and South American Bird Conservation Business Plan 2019.

Prem, M., Saavedra, S., Vargas, J.F., 2020. End-of-conflict deforestation: Evidence from Colombia’s peace agreement. World Dev. 129, 104852. https://doi.org/10.1016/j.worlddev.2019.104852

Presidencia de la Republica de Colombia, 2021. Aprueban $90 mil millones para proyectos ambientales y productivos en municipios PDET [WWW Document]. URL https://www.portalparalapaz.gov.co/publicaciones/1843/aprueban-90-mil-millones-para-proyectos-ambientales-y-productivos-en-municipios-pdet/ (accessed 3.30.22).

R Core Team, 2020. R: A language and environment for statistical computing.

Robinson, B.E., Masuda, Y.J., Kelly, A., Holland, M.B., Bedford, C., Childress, M., Fletschner, D., Game, E.T., Ginsburg, C., Hilhorst, T., Lawry, S., Miteva, D.A., Musengezi, J., Naughton-Treves, L., Nolte, C., Sunderlin, W.D., Veit, P., 2018. Incorporating Land Tenure Security into Conservation. Conserv. Lett. https://doi.org/10.1111/conl.12383

Rodríguez Eraso, N., Armenteras, D., Morales, M., Romero, M., 2016. Ecosistemas de los Andes Colombianos. Segunda edición. Bogota, Colombia.

Rosenberg, K. V., Dokter, A.M., Blancher, P.J., Sauer, J.R., Smith, A.C., Smith, P.A., Stanton, J.C., Panjabi, A., Helft, L., Parr, M., Marra, P.P., 2019. Decline of the North American avifauna. Science (80-.). 366, 120–124. https://doi.org/10.1126/science.aaw1313

RUNAP. Registro Único Nacional de Áreas Protegidas [WWW Document], 2021. URL https://runap.parquesnacionales.gov.co/ (accessed 3.1.21).

Runge, C.A., Watson, J.E.M., Butchart, S.H.M., Hanson, J.O., Possingham, H.P., Fuller, R.A., 2015. Protected areas and global conservation of migratory birds. Science (80-.). 350. https://doi.org/10.1126/science.aac9180

SARA. Species At Risk Act, 2002. Bill C-5, An act respecting the protection of wildlife species at risk in Canada.

Scarlett, L., McKinney, M., 2016. Connecting people and places: The emerging role of network governance in large landscape conservation. Front. Ecol. Environ. https://doi.org/10.1002/fee.1247

Schuster, R., Wilson, S., Rodewald, A.D., Arcese, P., Fink, D., Auer, T., Bennett, J.R., 2019. Optimizing the conservation of migratory species over their full annual cycle. Nat. Commun. 10. https://doi.org/10.1038/s41467-019-09723-8

Tejedor-Garavito, N., Álvarez, E., Arango-Caro, S., Araujo-Murakami, A., Blundo, C., Boza-Espinoza, T.E., La Torre-Cuadros, M.A., Gaviria, J., Gutiérrez, N., Jørgensen, P.M., León, B., López-Camacho, R., Malizia, L., Millán, B., Moraes, M., Pacheco, S., Rey-Benayas, J.M., Reynel, C., Timaná de la Flor, M., Ulloa-Ulloa, C., Vacas-Cruz, O., Newton, A.C., 2012. Evaluación del estado de conservación de los bosques montanos en los Andes tropicales. Ecosistemas 21, 148–166. https://doi.org/10.7818/re.2014.21-1-2.00

Torres Rodríguez, A.C., Binda, E., Ochoa Quintero, J.M., Garcia, H., Gómez, B., Soto, C., Martínez, S., Clerici, N., 2020. Answering the right questions. Addressing biodiversity conservation in post-conflict Colombia. Environ. Sci. Policy 104. https://doi.org/10.1016/j.envsci.2019.11.012

Vaast, P., Harmand, J.-M., Rapidel, B., Jagoret, P., Deheuvels, O., 2016. Coffee and Cocoa Production in Agroforestry—A Climate-Smart Agriculture Model, in: Climate Change and Agriculture Worldwide. https://doi.org/10.1007/978-94-017-7462-8_16

Vanegas Pinzón, S., Ospina Arango, O.L., Escobar Niño, G.A., Ramirez, W., Sánchez, J.J., 2015. Plan Nacional de Restauración: restauración ecológica, rehabilitación y recuperación de áreas disturbadas. Bogotá, Colombia.

Vélez, D., Tamayo, E., Ayerbe-Quiñones, F., Torres, J., Rey, J., Castro-Moreno, C., Ramírez, B., Ochoa-Quintero, J.M., 2021. Distribution of birds in Colombia. Biodivers. Data J. 9. https://doi.org/10.3897/BDJ.9.e59202

Waldron, A., Garrity, D., Malhi, Y., Girardin, C., Miller, D.C., Seddon, N., 2017. Agroforestry Can Enhance Food Security While Meeting Other Sustainable Development Goals. Trop. Conserv. Sci. 10. https://doi.org/10.1177/1940082917720667

Wilson, S., Lin, H.Y., Schuster, R., González, A.M., Gómez, C., Botero-Delgadillo, E., Bayly, N.J., Bennett, J.R., Rodewald, A.D., Roehrdanz, P.R., Ruiz Gutierrez, V., 2022. Opportunities for the conservation of migratory birds to benefit threatened resident vertebrates in the Neotropics. J. Appl. Ecol. 59, 653–663. https://doi.org/10.1111/1365-2664.14077

Wilson, S., Saracco, J.F., Krikun, R., Flockhart, D.T.T., Godwin, C., Foster, K.R., 2018. Drivers of demographic decline across the annual cycle of a threatened migratory bird. Sci. Rep. 8. https://doi.org/10.1038/s41598-018-25633-z

Wilson, S., Schuster, R., Rodewald, A.D., Bennett, J.R., Smith, A.C., La Sorte, F.A., Verburg, P.H., Arcese, P., 2019. Prioritize diversity or declining species? Trade-offs and synergies in spatial planning for the conservation of migratory birds in the face of land cover change. Biol. Conserv. 239, 108285. https://doi.org/10.1016/j.biocon.2019.108285

Worsdell, T., Kumar, K., Allan, J.R., Gibbon, G.E.M., White, A., Khare, A., Frechette, A., 2020. Rights-Based Conservation: The path to preserving Earth’s biological and cultural diversity? Washington, DC.

Zúñiga-Upegui, P., Arnaiz-Schmitz, C., Herrero-Jáuregui, C., Smart, S.M., López-Santiago, C.A., Schmitz, M.F., 2019. Exploring social-ecological systems in the transition from war to peace: A scenario-based approach to forecasting the post-conflict landscape in a Colombian region. Sci. Total Environ. 695. https://doi.org/10.1016/j.scitotenv.2019.133874

