## Supplemental Material for "Habitat protection and restoration: win-win opportunities for migratory birds in the Northern Andes"

Supplemental Material Table 1. Direct causes of deforestation in the Colombian Andes from 2010-2015, agents and indirect causes of deforestation. *Direct causes and agents operating below 1000 m asl. Data summarized from (González et al., 2018).

| Direct Causes and agents | **Indirect causes |
|---|---|
| **Traditional agricultural production** | 1-7 |
| Producers using traditional cropping techniques | |
| **Palm oil, industrial agricultural production*** | |
| Industrial crop producers* | |
| **Coca production** | 1,2,4,8-11 |
| Coca producers | |
| **Artisanal coal extraction*** | |
| Formal miner* | |
| Informal miner* | |
| **Infrastructure expansion (Transportation)*** | |
| Formal constructor* | |
| Informal constructor* | |
| **Logging** | 1,4,6,12-13 |
| Informal logger (timber to sell) | |
| Informal logger (timber for consumption) | |
| **Cattle ranching** | 3,5,9,14-19 |
| Large-scale rancher | |
| **Open-pit gold extraction** | 1,6,9,11,20-22 |
| Formal miner | |
| Informal Miner | |
| **Open-pit gold extraction and other minerals** | 1,6,9,11,20-22 |
| Formal miner | |
| Informal Miner | |

**Indirect causes driving the agents' decisions to transform the forest. 1. Poverty, 2. Limited access to productive technologies, 3. Unsustainable traditional practices, 4. Low state presence, 5. High national market demand, 6. Unemployment, 7. Absence of fiscal policy that promotes effective use of land in rural areas, 8. International cocaine price, 9. Illegal economies, 10. Economical support expectations, 11. Armed conflict, 12. Lack of forestry policies for forest management, 13. Timber demand for productive activities, 14. Poor production technologies, 15. Inconsistent agricultural policy, 16. Low production cost, 17. Government incentives, 18. Political interest conflicts, 19. Territorial control, 20. International gold price, 21. Government support, 22. Low government control.

Supplemental Material Table 2. Protected areas classification according to the International Union for Conservation of Nature (IUCN) and their equivalent according to the Colombian National Protected Areas System (SINAP "Sistema Nacional de Areas Protegidas). Protected areas are managed at the national government level by the National Natural Park System, and the regional government level by Autonomous Regional Corporations (CARs), and by private landowners.

| IUCN Category | Protected Area Category (SINAP) | Management Agency |
|---|---|---|
| (Ib) Wilderness Areas | Fauna and Flora Sanctuaries | National Natural Park System |
| | Flora Sanctuaries | National Natural Park System |
| (II) National Parks | Regional Natural Parks | CARs |
| | National Natural Parks | National Natural Park System |
| (V) Protected Landscapes | Recreational Areas | CARs |
| (VI) Protected Areas with Sustainable Use of Natural Resources | National Forest Reserves | National Natural Park System |
| | Regional Forest Reserves | CARs |
| | Regional Integrated Management Districts | CARs |
| | Soil Conservation Districts | CARs |
| | Civil Society Natural Reserves | Private lands |

Supplemental Material Table 3. Land-use across Migrant Focal Areas according to the CORINE land use-land cover dataset categories.

| Label | Land use classification | Area (ha) |
|---|---|---|
| Dense forest | Forest | 3,482,625 |
| Open forest | Forest | 16,062 |
| Fragmented forest | Forest | 386,967 |
| Riparian forest | Forest | 229,714 |
| Other temporary crops | Agricultural areas_crops_mosaic | 285 |
| Cereal crops | Agricultural areas_crops_mosaic | 2 |
| Vegetables | Agricultural areas_crops_mosaic | 126 |
| Tubers | Agricultural areas_crops_mosaic | 94 |
| Permanent arable crops | Agricultural areas_crops_mosaic | 27,647 |
| Confined crops (tree nurseries) | Agricultural areas_crops_mosaic | 432 |
| Mosaic of crops | Agricultural areas_crops_mosaic | 56,047 |
| Mosaic of pastures and crops | Agricultural areas_crops_mosaic | 836,365 |
| Mosaic of crops, pastures and natural spaces | Agricultural areas_crops_mosaic | 1,312,155 |
| Mosaic of pastures with natural spaces | Agricultural areas_crops_mosaic | 741,122 |
| Mosaic of crops with natural spaces | Agricultural areas_crops_mosaic | 327,929 |
| Bushy permanent crops | Agroforesty areas with tree or shurbs and forest plantation | 254,215 |
| Agroforestry | Agroforesty areas with tree or shurbs and forest plantation | 82,507 |
| Pastureland without trees | Agricultural areas pastures | 1,464,012 |
| Pastures with trees | Agricultural areas_pastures | 18,942 |
| Pastures and weed | Agricultural areas_pastures | 307,344 |
| Grasslands | Early succession stage | 100,579 |
| Shrubs | Early succession stage | 89,948 |
| Secondary vegetation or in transition | Early succession stage | 1,096,353 |
| Sandy natural areas | Not included | |
| Rocky outcrop | Not included | |
| Uncovered degraded lands | Not included | |
| Burnt areas | Not included | |
| Wetlands | Not included | |
| Rivers (50m) | Not included | |
| Lagoons, lakes and y Cienegas | Not included | |
| Channels | Not included | |
| Artificial waterbodies | Not included | |
| Continuous urban areas | Not included | |
| Discontinuous urban areas | Not included | |
| Industrial and commercial zones | Not included | |

| | | |
|---|---|---|
| Roads, rail freight terminals and associated terrains. | Not included | |
| Airports | Not included | |
| Hydraulic plants | Not included | |
| Mining areas | Not included | |
| Garbage dumps | Not included | |
| Urban parks | Not included | |
| Recreational areas | Not included | |
| | Total | 10,831,472 |

Supplemental Material Table 4. Area and % of land of Regional Autonomous Corporations (CARs) within Forest, Restoration and Rehabilitation Planning Areas. CARs with the largest amount of land and adding 50% of the area in each planning area are shown in bold. Total %= Percentage of Forested, Restoration and Rehabilitation Areas within the jurisdiction of the CARs.

| Regional Autonomous Corporation | Forested Areas | % | Restoration Areas | % | Rehabilitation Areas | % |
|---|---|---|---|---|---|---|
| CORPOAMAZONIA | **383267** | **9.31** | 3570 | 1.30 | 1855 | 0.59 |
| CORPONARIÑO | **337500** | **8.20** | 3271 | 1.19 | 2594 | 0.83 |
| CRC | **287410** | **6.98** | **15564** | **5.65** | 27313 | 8.71 |
| CODECHOCO | **280315** | **6.81** | 179 | 0.06 | 1134 | 0.36 |
| CORPORINOQUIA | **241875** | **5.88** | 3911 | 1.42 | 2923 | 0.93 |
| CORMACARENA | **218029** | **5.30** | 764 | 0.28 | 931 | 0.30 |
| CORPONOR | 197805 | 4.81 | **17154** | **6.22** | 11440 | 3.65 |
| CAM | 188216 | 4.57 | **34877** | **12.65** | **52322** | **16.69** |
| CORPOURABA | 172160 | 4.18 | 4601 | 1.67 | 4627 | 1.48 |
| CORANTIOQUIA | 171757 | 4.17 | 11817 | 4.29 | 18875 | 6.02 |
| CSB | 122299 | 2.97 | 7613 | 2.76 | 4896 | 1.56 |
| CVC | 120209 | 2.92 | **15681** | **5.69** | 14666 | 4.68 |
| CAS | 107847 | 2.62 | **33959** | **12.32** | **41505** | **13.24** |
| CORTOLIMA | 97243 | 2.36 | 14072 | 5.10 | **30135** | **9.61** |
| CORNARE | 86347 | 2.10 | 5784 | 2.10 | 5554 | 1.77 |
| CARDER | 58965 | 1.43 | 7777 | 2.82 | 3902 | 1.24 |
| CORPOBOYACA | 51252 | 1.25 | 2152 | 0.78 | 2238 | 0.71 |
| CORPOCESAR | 39166 | 0.95 | 5576 | 2.02 | 7164 | 2.29 |
| CORPOGUAVIO | 33228 | 0.81 | 1247 | 0.45 | 1899 | 0.61 |
| CORPOCALDAS | 31656 | 0.77 | 1871 | 0.68 | 3804 | 1.21 |
| CDMB | 27826 | 0.68 | 9771 | 3.54 | 4799 | 1.53 |
| CAR | 25472 | 0.62 | **33047** | **11.99** | **48190** | **15.37** |
| CORPOCHIVOR | 23901 | 0.58 | 2213 | 0.80 | 3424 | 1.09 |
| CRQ | 10427 | 0.25 | 5884 | 2.13 | 1505 | 0.48 |
| CVS | 8090 | 0.20 | 1800 | 0.65 | 2427 | 0.77 |
| Total | 3322262 | 80.1 | 244155 | 88.6 | 300122 | 95.8 |

**CORPOAMAZONIA**= Corporación para el Desarrollo Sostenible del Sur de la Amazonia, **CORPONARIÑO**= Corporación Autónoma Regional de Nariño**, CRC** = Corporación Autónoma Regional del Cauca, **CODECHOCO**= Corporación Autónoma Regional para el Desarrollo Sostenible de Choco, **CORPORINOQUIA**= Corporación Autónoma Regional de la Orinoquia, **CORMACARENA**= Corporación para el Desarrollo Sostenible del Área de Manejo Especial La Macarena**, CORPONOR**=Corporación Autónoma Regional de la Frontera Nororiental, **CAM**=

Corporación Autónoma Regional del Alto Magdalena, **CORPOURABA**= Corporación para el Desarrollo Sostenible del Urabá, **CORANTIOQUIA**= Corporación Autónoma Regional del Centro de Antioquia, **CSB**= Corporación Autónoma Regional del Sur de Bolívar, **CVC**= Corporación Autónoma Regional del Valle del Cauca, **CAS**= Corporación Autónoma Regional de Santander, **CORTOLIMA**= Corporación Autónoma Regional del Tolima, **CORNARE**= Corporación Autónoma Regional de las Cuencas de los Ríos Negro y Nare, **CARDER**= Corporación Autónoma Regional de Risaralda, **CORPOBOYACA**= Corporación Autónoma Regional de Boyacá, **CORPOCESAR**= Corporación Autónoma Regional del Cesar, **CORPOGUAVIO**= Corporación Autónoma Regional del Guavio, **CORPOCALDAS**= Corporación Autónoma Regional de Caldas, **CDMB**= Corporación Autónoma Regional para la Defensa de la Meseta de Bucaramanga, **CAR**= Corporación Autónoma Regional de Cundinamarca, **CORPOCHIVOR**= Corporación Autónoma Regional de Chivor, **CRQ**= Corporación Autónoma Regional del Quindío, **CVS**= Corporación Autónoma Regional de los Valles del Sinú y del San Jorge.

Supplemental Material Figure 1. Autonomous Regional Corporations (CARs) with ≥50% of forested areas within their jurisdictions: Corpoamazonia (383,267 ha), Corponariño (337,500 ha), Corporación Autónoma Regional del Cauca (287,410 ha), Codechoco (280,315 ha), Corporinoquia (241,875 ha), and Cormacarena (241,875 ha). Forested areas = 4,115,368 ha; Forested Area within CARs's jurisdictions= 3,322,262 ha (81%).

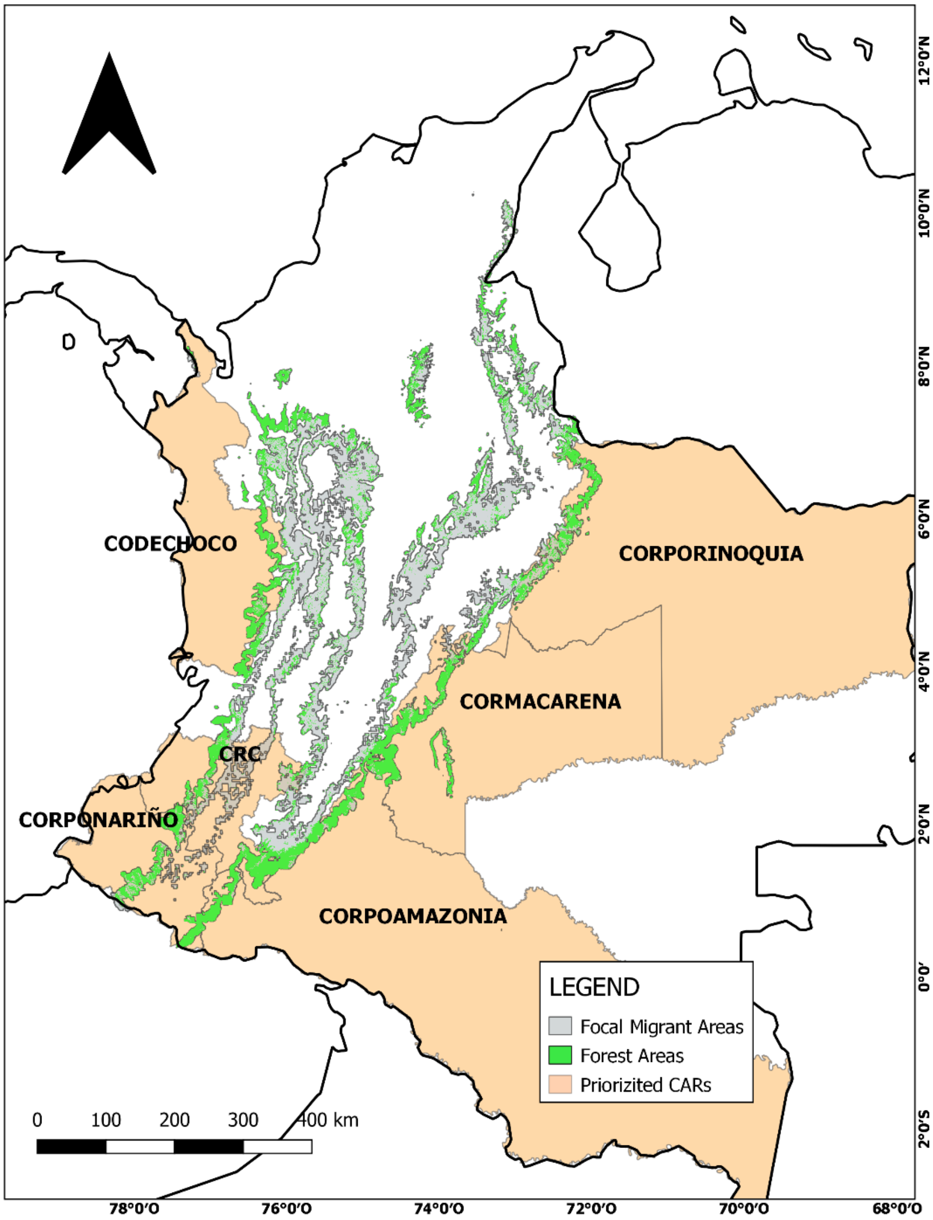

Supplemental Material Figure 2. Autonomous Regional Corporations (CARs) with ≥50% of restoration and rehabilitation areas within their jurisdictions. CRC= Corporación Autónoma Regional del Cauca (3,090,911 ha), CORTOLIMA= Corporación Autónoma Regional del Tolima (2,415,020 ha), CAS= Corporación Autónoma Regional de Santander (2,593,983 ha), CAR= Corporación Autónoma Regional de Cundinamarca (1,827,952 ha), CAM= Corporación Autónoma Regional del Alto Magdalena (1,813,533 ha).

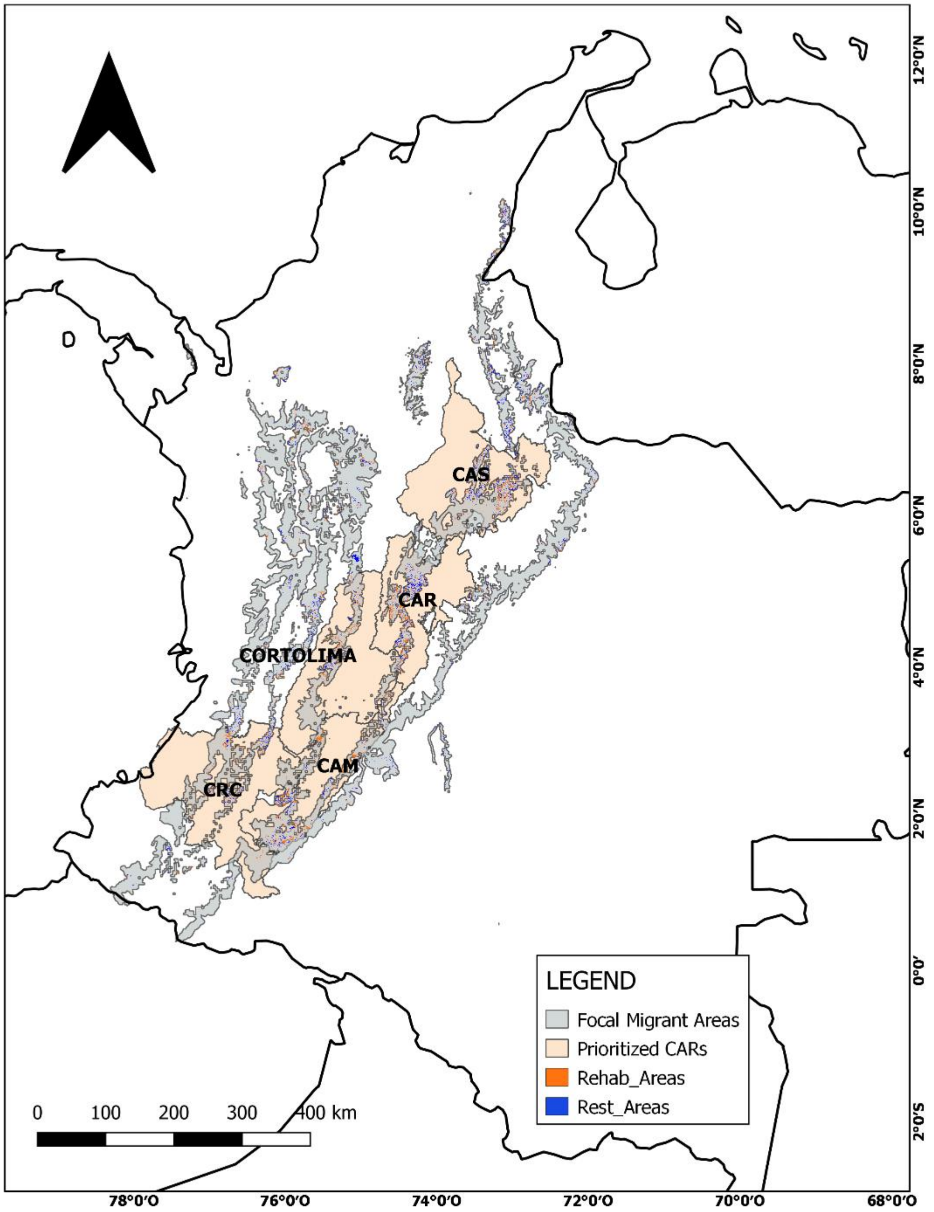
